# Reconstruction of the Hypothalamo-Neurohypophysial System and Functional Dissection of Magnocellular Oxytocin Neurons in the Brain

**DOI:** 10.1101/2020.03.26.007070

**Authors:** Bin Zhang, Liyao Qiu, Wei Xiao, Hong Ni, Lunhao Chen, Fan Wang, Weihao Mai, Hui Gong, Shumin Duan, Anan Li, Zhihua Gao

## Abstract

The hypothalamo-neurohypophysial system (HNS), comprising hypothalamic magnocellular neuroendocrine cells (MNCs) and the neurohypophysis, plays a pivotal role in regulating reproduction and fluid homeostasis by releasing oxytocin and vasopressin into the bloodstream. However, it remains incompletely understood on its structure and whether it contributes to the central actions of oxytocin and vasopressin. Using viral tracing and whole brain imaging, we reconstructed the three-dimensional architecture of the HNS and uncovered that subsets of MNCs collaterally project to multiple extrahypothalamic regions. Moreover, selective activation of magnocellular oxytocin neurons promoted peripheral oxytocin release and facilitated central oxytocin-mediated social interactions. Further, MNCs-released oxytocin in the caudate putamen enhanced locomotion to orchestrate social investigation. Our work reveals the previously unrecognized complexity of the HNS and provides structural and functional evidence for MNCs in coordinating both peripheral and central oxytocin-mediated actions, which will shed light on the mechanistic understanding of oxytocin-related psychiatric diseases.

## INTRODUCTION

The hypothalamo-neurohypophysial system (HNS), comprising hypothalamic magnocellular neuroendocrine cells (MNCs) and their axonal projections to the posterior pituitary (PPi), is a key gateway for the brain to regulate peripheral function. Primarily residing in the paraventricular nucleus (PVN) and supraoptic nucleus (SON), MNCs release oxytocin (OXT) and arginine vasopressin (AVP) from their neurohypophysial terminals into the peripheral bloodstream to regulate reproduction and fluid homeostasis (Harris, 1955b; Insel et al., 1997; Leng et al., 2015). Despite the pivotal importance of the HNS in neuroendocrine regulation, knowledge on the structure of the HNS is mainly from histological tract tracing studies conducted decades ago (Buijs et al., 1978; Hou-Yu et al., 1986; Sofroniew and Glasmann, 1981). The deep location of the pituitary and the presence of the fragile neural stalk leave the system frequently overlooked in recent brain-wide connectome studies (Dong 2008, Oh, Harris et al. 2014). To date, the full composition and wiring of the HNS remains incompletely understood.

On the other hand, accumulating evidence demonstrate that central OXT and AVP are involved in the regulation of various social and stress related behaviors in the brain (Grinevich and Stoop, 2018; Jurek and Neumann, 2018; Onaka et al., 2012; Ross and Young, 2009). Moreover, defects in the oxytocinergic system is found associated with mental diseases such as autism and anxiety-related disorders (Andari et al., 2010; Hollander et al., 2007; Windle et al., 1997). MNCs are known to send unipolar axons to the neurohypophysis and peripheral peptides rarely cross the blood-brain barrier (Armstrong and Paxinos, 1995; Armstrong et al., 1980; Leng and Ludwig, 2016; Mens et al., 1983; Swanson and Kuypers, 1980); however, whether and how they contribute to central actions remains controversial.

Other than the PPi-projecting magnocellular OXT (Magno-OXT) neurons (20-35 µm soma diameter), PVN also contains centrally-projecting parvocellular OXT (Parvo-OXT) neurons (10-15 µm soma diameter), which were thought to be responsible for the central actions of OXT (Badoer, 1996; Insel et al., 1997; Lee et al., 2009; Swanson and Kuypers, 1980). Parvo-OXT neurons contain much less OXT than Magno-OXT neurons and mainly project to the hindbrain and spinal cord (Swanson and Kuypers, 1980; Swanson and Sawchenko, 1980, 1983); however, whether they participate in the OXT-mediated actions in the forebrain remains elusive (Dolen et al., 2013). Somatodendritic release from Magno-OXT neurons was the main source of OXT in the brain, and was involved in central effects of OXT such as social recognition and maternal behavior (Bergquist and Ludwig, 2008; Ludwig and Leng, 2006; Takayanagi et al., 2017). Given the limited diffusion of locally-released peptides to a distant region (Chini et al., 2017), it is less likely that the central actions of OXT observed in multiple extrahypothalamic regions were effects of locally-released OXT (Li et al., 2016; Marlin et al., 2015; Nakajima et al., 2014).

Using OXT promoter-driven adeno-associated viruses (AAV) that specifically infect OXT neurons, recent studies found that OXT neurons extensively projected to more than 50 brain regions and that axonal release of OXT neurons contributed to OXT-mediated fear attenuation (Knobloch et al., 2012; Menon et al., 2018). In these studies, however, viruses were injected into the hypothalamus and thus could infect both Magno-OXT and Parvo-OXT neurons (Knobloch et al., 2012). As such, the processes and/or fibers observed within the brain may not necessarily represent projections of Magno-OXT neurons. Instead, different subsets of OXT neurons may independently project to different targets, confounding the origins of OXT in the brain. Mapping the connectivity of MNCs, particularly Magno-OXT neurons in the HNS, is therefore crucial to address these concerns.

As all MNCs project to the neurohypophysis, we reasoned that retrograde tracing from the PPi is a proper way to selectively label MNCs. Together with newly engineered viral tracers, we may be able to fully delineate the morphology and projections of MNCs. By injecting the green fluorescence protein (GFP)-tagged retrogradely transported virus (rAAV2-Retro-GFP) into PPi (Tervo et al., 2016), we selectively labeled the MNC ensemble and reconstructed the three-dimensional (3D) connection map of the rat HNS. We further traced their fiber trajectory at single-cell resolution and uncovered multiple extrahypothalamic collateral projections of MNCs. Using customized OXT-Cre rats, we verified that Magno-OXT neurons collaterally project to multiple areas, including the previously unidentified caudate putamen (CPu). Chemogenetic activation of Magno-OXT neurons not only elevates peripheral OXT release but also drives central OXT-dependent social interactions and locomotion. This study uncovers the full organization of the HNS and provides direct evidence for Magno-OXT neurons in coordinating both central and peripheral actions.

## RESULTS

### Retrograde Viral Tracing from the PPi Labeled the MNC Ensemble

To selectively label the MNCs projecting to the neurohypophysis, we injected the retrogradely-transported virus rAAV2-Retro-GFP (Retro-GFP) into the PPi of SD rats (Figure 1A Left). After two weeks, intense GFP epifluorescence was observed in the PPi, demonstrating the accuracy of viral injection and infection (Figure 1A Middle). Robust GFP signals within the neural stalk reflected the effective retrograde transport of the virus along the axons (Figure 1A Right).

**Figure 1.**
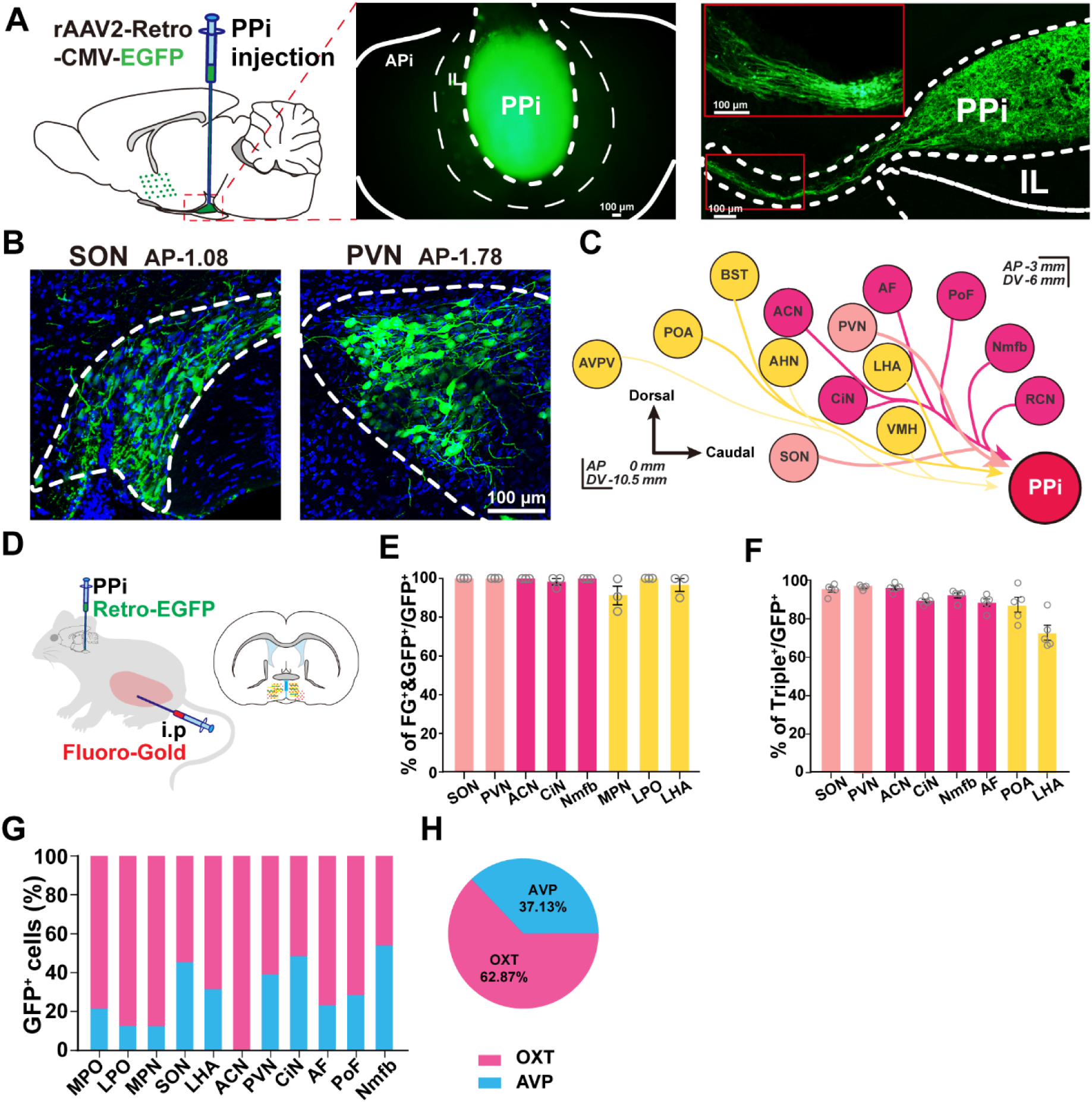
Retrograde tracing of magnocellular neuroendocrine cells (MNCs) in multiple hypothalamic regions projecting to the neurohypophysis. (A) Schematic injection and viral infection of rAAV2-Retro-CMV-GFP in the posterior pituitary (PPi). Middle and Right, a representative image of the viral infection in the PPi and the retrograde transport within the pituitary stalk from the PPi. (B) Representative images of PPi-retrogradely traced (GFP^+^) cells in SON and PVN. (C) A diagram summarizing the connections between hypothalamus and neurohypophysis. The hypothalamic nuclei were assembled relative to their rostral/caudal and dorsal/ventral locations and divided into three subpopulations. The light pink, deep pink and yellow circles represent the principal, accessory and scattered magnocellular neuroendocrine system (PMN, AMN and SMN), respectively. (D) Schematic representation of the Fluoro-Gold (FG) injection 14 days after rAAV2-Retro-GFP labeling. (E) Histogram of the percentage of FG^+^ cells in retrogradely traced cells (GFP^+^) (n=3 rats). (F) Statistical analysis of AVP^+^ or OXT^+^ neurons in retrogradely traced cells (GFP^+^) in different nuclei (n=5 rats). (G) Percentage of AVP^+^ or OXT^+^ cells within different magnocellular nuclei (n=5 rats). (H) A pie chart showing the ratio of AVP^+^ and OXT^+^ GFP-positive neurons in all the traced MNCs (n=5 rats). AP, anterior-posterior distance from the Bregma (mm). Scale bar: 100 µm.

GFP^+^ cells were primarily located from the preoptic to the tuberal areas of the hypothalamus and packed into multiple discrete nuclei, ranging from 0 mm to −3 mm along the anterior-posterior (AP) axis and −6 mm to −10.5 mm along the dorsal-ventral (DV) axis (Figure 1C) in both male and female rats. Abundant GFP^+^ cells were distributed in the expected principal magnocellular neurosecretory nuclei (PMN), SON and PVN (Figure 1B). Moreover, densely packed GFP^+^ cells were seen in the less-defined accessory magnocellular neurosecretory nuclei (AMN) (Fisher et al., 1979; Peterson, 1966; Rhodes et al., 1981), consisting of the anterior commissural nucleus (ACN), the nucleus circularis (CiN), the anterior perifornical nucleus (AF), the posterior perifornical nucleus (PoF), the nucleus of the medial forebrain bundle (Nmfb), and the retrochiasmatic nucleus (RCN) (Figure S1A and S1B). Notably, scattered GFP^+^ cells were also seen in the lateral and preoptic hypothalamic areas, including the bed nucleus of the stria terminalis (BST), medial preoptic nucleus (MPN), medial preoptic area (MPO), and lateral preoptic area (LPO) (Figure S1C), as well as the anteroventral periventricular nucleus (AVPV) and preoptic periventricular nucleus (PVpo) (Figure S1B and data not shown). These areas are herein referred to as the scattered magnocellular neurosecretory system (SMN). Thus, the engineered retrograde viral tracers allowed us to efficiently label the cell ensemble that directly innervates the neurohypophysis from multiple nuclei of the hypothalamus (schematically illustrated in Figure 1C and Figure S1C).

### Retrogradely-traced MNCs Project beyond the Blood-brain-barrier and Contain AVP and/or OXT

MNCs release hormones directly into the bloodstream via their axonal terminals in the neurohypophysis devoid of the blood-brain-barrier (BBB). To assess the neuroendocrine property of the virally-traced GFP^+^ cells, we intraperitoneally injected the rats (PPi-viral injected) with Fluoro-Gold (FG), a fluorescent dye that can be taken up by nerve terminals outside the BBB and retrogradely transported to the somas (Figure 1D). Prominent FG-labeled neurons were observed in the SON, PVN and arcuate nucleus that are known to be neurosecretory (Figure S2A and data not shown for ARC). Importantly, all the GFP^+^ cells in the SON, PVN and AMN were co-labeled with FG (100% dual labeling). GFP^+^ cells scattered in the preoptic area and lateral hypothalamic area also exhibited a high percentage (89% and 95%, respectively) of FG dual-labelling (Figure 1E), suggesting that retrograde tracing from the PPi faithfully labelled the MNCs innervating the PPi. Given the limited efficiency of viral infection and retrograde transport, FG^+^ MNCs within the SON, PVN and AMN were partly labeled by GFP. Approximately 61% of FG^+^ cells in SON were GFP positive (Figure S2B), which reflects the labelling efficiency of the retrograde viral tracer, as virtually all neurons in the rat SON project to the neurohypophysis (Swanson and Sawchenko, 1983).

MNCs are known to contain AVP/OXT. To examine whether all traced MNCs contain AVP or OXT, we performed co-immunostaining of GFP with mixed antibodies to vasopressin and oxytocin (antibodies were both generated from rabbits). As expected, almost all GFP^+^ cells in the PMNs and AMNs were co-labeled with AVP and OXT (Figure 1F and Figure S3A). By contrast, only 80% of GFP^+^ cells in LHA expressed AVP or OXT, suggesting that not all cells projecting to the neurohypophysis contain AVP or OXT. Using OXT and AVP antibodies raised from different species, we were able to determine the fraction of OXT or AVP cells in different nuclei. Consistent with previous findings (Hou-Yu et al., 1986; Sofroniew, 1983), we found that ACN contained only OXT neurons, whereas other nuclei contained both AVP and OXT with different percentages (Figure S3B). All the nuclei, except the Nmfb, contained more OXT neurons than AVP neurons (Figure 1G). Altogether, OXT^+^ and AVP^+^ cells accounted for 62% and 37% of retrogradely-traced MNCs (Figure 1H).

### 3D Reconstruction of the HNS by Fluorescent Micro-Optical Sectioning Tomography (fMOST)

Given that retrograde viral tracer clearly labeled the MNC ensemble, we took a further step to construct the HNS. To acquire a full architecture of the HNS, the brains with intact neural stalks and pituitaries were carefully isolated from PPi retro-traced rats and imaged by fMOST at a resolution of 0.32×0.32×2 µm^3^ (schematically illustrated in Figure 2A). Individual dataset containing over 6600 coronal slices from each brain was generated and the raw data were registered to the rat brain atlas (Swanson, 2004). Since MNCs were localized between 0 and −3.4 mm along the AP axis, we axially stacked 850 re-sampled fMOST coronal images spanning this area to reconstruct the HNS using Imaris software (Figure 2B and 2C). The reconstructed projection gives a three-dimensional (3D) overview of the HNS, with MNCs (5868 cells/rat on average) packed into more than 8 nuclei in the hypothalamus and axons bundled together to form the arc tracts of Greving (Greving, 1926), subsequently passing through the median eminence and terminating in the posterior pituitary (Figure 2B). Among the multiple nuclei, SON, abutting the ventral surface of the brain (from 0 to −1.8 mm along the AP axis), contains the most abundant GFP^+^ cells (1617±198 cells). Caudal and medial to SON lies the RCN, whereas dorsal to the SON is the Nmfb. Notably, MNCs in the PVN and ACN are arranged into a butterfly-like shape, with ACN constituting the forewings and PVN forming the hindwings. The perifornical MNC aggregates, including both AF and PoF, lie dorsal-lateral to the fornix, whereas CiN lies midway between the PVN and SON (Figure 2B and 2C).

**Figure 2.**
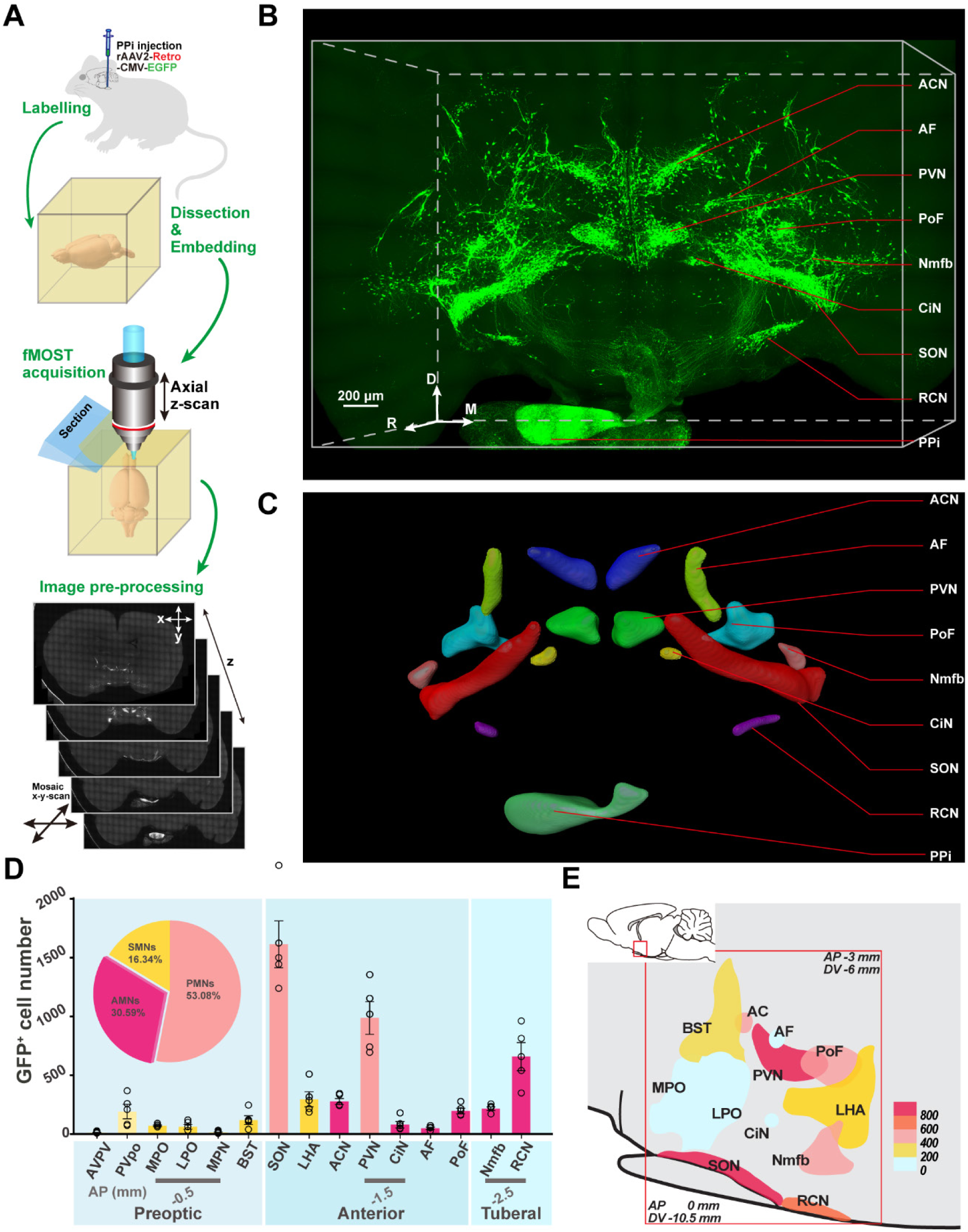
3D reconstruction of the HNS using fluorescent Micro-Optical Sectioning Tomography (fMOST) (A) Schematic procedures of the fMOST imaging. (B and C) Three-dimensional projections of the HNS derived from 1700 fMOST images (thickness of 3400 µm). A coronal view of the 3D HNS (B) and the multi-color registration of the nuclei (C). (D) Quantitative analysis of GFP^+^ cells in different nuclei indicating distinctive subpopulations: (PMNs, light pink), AMNs (deep pink), and the scattered MNs (yellow) (n=5 rats). The pie chart shows the percentage of GFP^+^ neurons in PMNs, AMNs and SMNs (n=5 rats). (E) A heatmap demonstrating the distribution of GFP^+^ neurons, located from 0 mm to −3 mm in AP axis and from −6 mm to −10.5 mm in DV axis. The number of cells in individual areas is color-coded as indicated. n=5 rats. C, caudal; D, dorsal; M, medial; R, rostral. Scale bar: 200 μm.

Three-dimensional measurement of the MNCs reveals that the average volume of GFP^+^ cells is above 1300 µm^3^ (approximately 25 µm in diameter), which meets the criteria for the size of MNCs (20-35 µm) (Figure S4A and S4B) (Sofroniew and Glasmann, 1981; Stern and Armstrong, 1998). Quantitatively, about half (53±2.7%) of the GFP^+^ MNCs were located in the PMN including SON and PVN, and one third (31±2.3 %) and 16% of labeled MNCs were found in the AMN and SMN, respectively (Figure 2D). A heatmap illustrating the number and distribution of GFP^+^ MNCs in different nuclei is presented on a sagittal plane of the rat brain (Figure 2E).

### Collateral Projections of MNCs within the Brain

High-resolution fMOST images allowed us to identify not only the somas but also the fine processes of MNCs (Figure 3A). Within the hypothalamus, the Greving tract of MNC axons was clearly seen (Figure 2B); outside the hypothalamus, GFP^+^ processes/fibers were also observed in multiple areas. Both the cortical regions, such as the piriform cortex (Piri C) and auditory cortex (Audi C), and the subcortical regions, including the amygdala (AMY), nucleus accumbens (NAc), caudate putamen (CPu) and lateral septum (LS), contained GFP^+^ fibers (Figure 3B and S4C), with peak density of fibers found in the piriform cortex (Figure 3C and 3D). Of note, GFP^+^ fibers were undetectable in the reward center, ventral tegmental area (VTA) (Figure 3B), in accordance with recent findings showing that Magno-OXT neurons do not project to the VTA in mice (Xiao et al., 2017). All the GFP^+^ areas, except amygdala, are more than 1000 µm away from the somal location of MNCs in the hypothalamus. As the viruses were injected into the PPi, fibers in the extra-hypothalamic regions likely represent the collateral projections of the retrogradely traced MNCs in the brain, as schematically summarized in Figure 3E.

**Figure 3.**
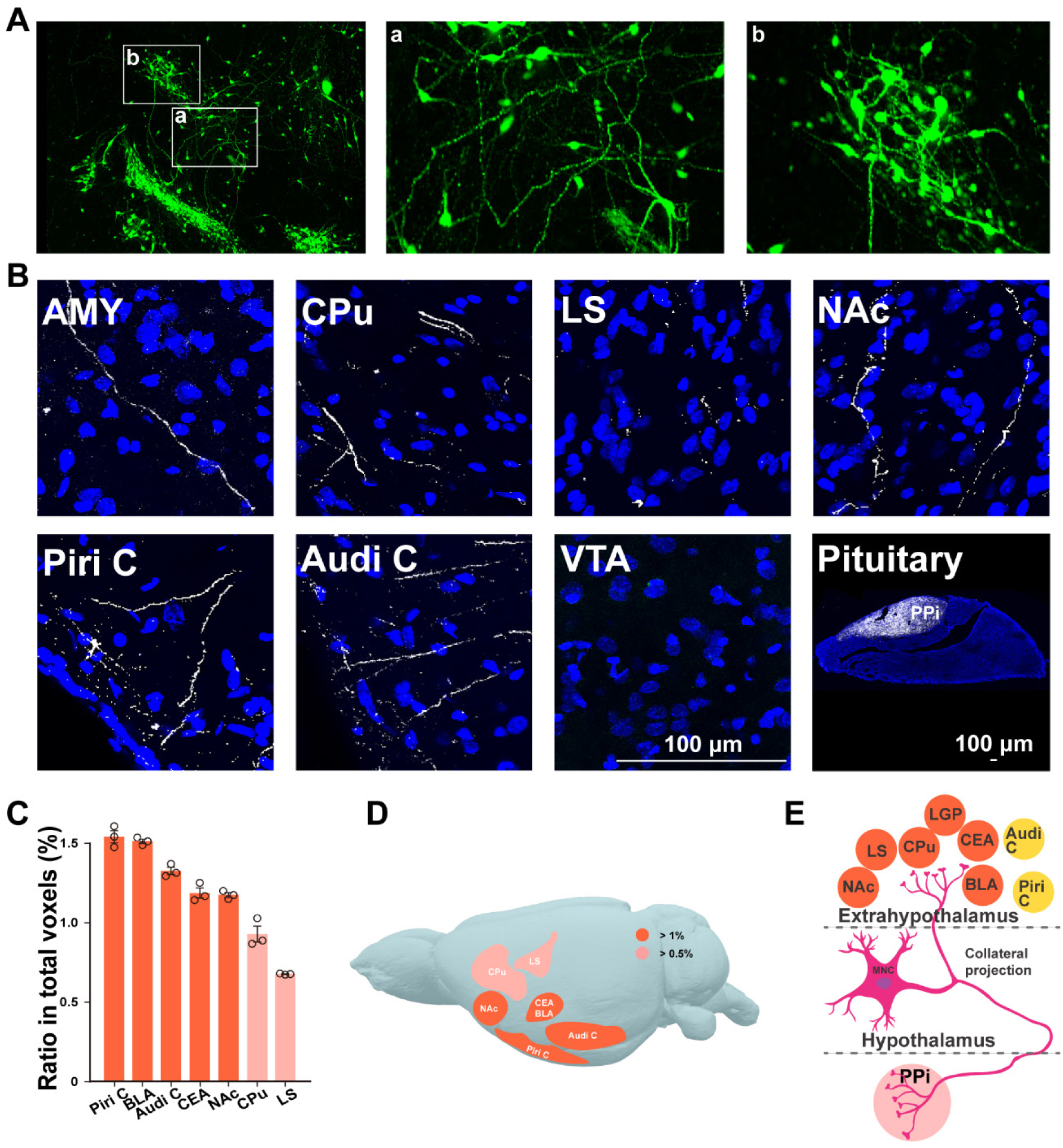
Processes of retrogradely-labeled MNCs in cortical and subcortical regions. (A) Representative fMOST images showing the processes (a) and somas (b) of PPi-retrogradely traced cells. (B) Representative images of GFP^+^ fibers in extra-hypothalamic areas including amygdala (AMY), caudate putamen (CPu), lateral septum (LS), piriform cortex (Piri C), auditory cortex (Audi C), ventral tagmental area (VTA) and the posterior pituitary (PPi) (C and D) Histogram (C) and heatmap (D) of collateral distribution in cortical and subcortical regions. The density of GFP^+^ fibers in extra-hypothalamic areas was calculated by the ratio of voxels containing GFP to total voxels in individual nucleus (n=3 rats). (E) Schematic illustration of an MNC with collateral projections to posterior pituitary and extra-hypothalamus. Scale bar: 100 μm.

To fully delineate the morphology of MNCs, we traced fiber trajectories of GFP-labeled MNCs and reconstructed the cells at single-cell resolution (Figure 4A, 4B, S4D and S4E). All the 46 traced MNCs send axons winding through the anterior/lateral hypothalamic area (AHA/LHA), passing by the fornix and the ventromedial hypothalamic nucleus (VMH), to the medial eminence (ME) (Figure 4B and 4C). Other than ME projection, 7 cells (7/46, 15%) within PVN and PoF showed bifurcated projections to the NAc, lateral globus pallidus (LGP) and CPu (Figure 4D). Injection of the retrograde viral tracer expressing Cre recombinase (rAAV2-Retro-Cre) into the PPi and a Cre-dependent AAV expressing GFP and synaptophysin-conjugated mRuby into the PVN (Figure 4E) revealed prominent GFP signals and mRuby-positive inflated beaded axon terminals in both the shell and core region of the NAc (Figure 4F and 4G), verifying the collateral projections to NAc from MNCs. Together, these data indicate that the PPi-projecting MNCs collaterally projected to multiple extrahypothalamic areas.

**Figure 4.**
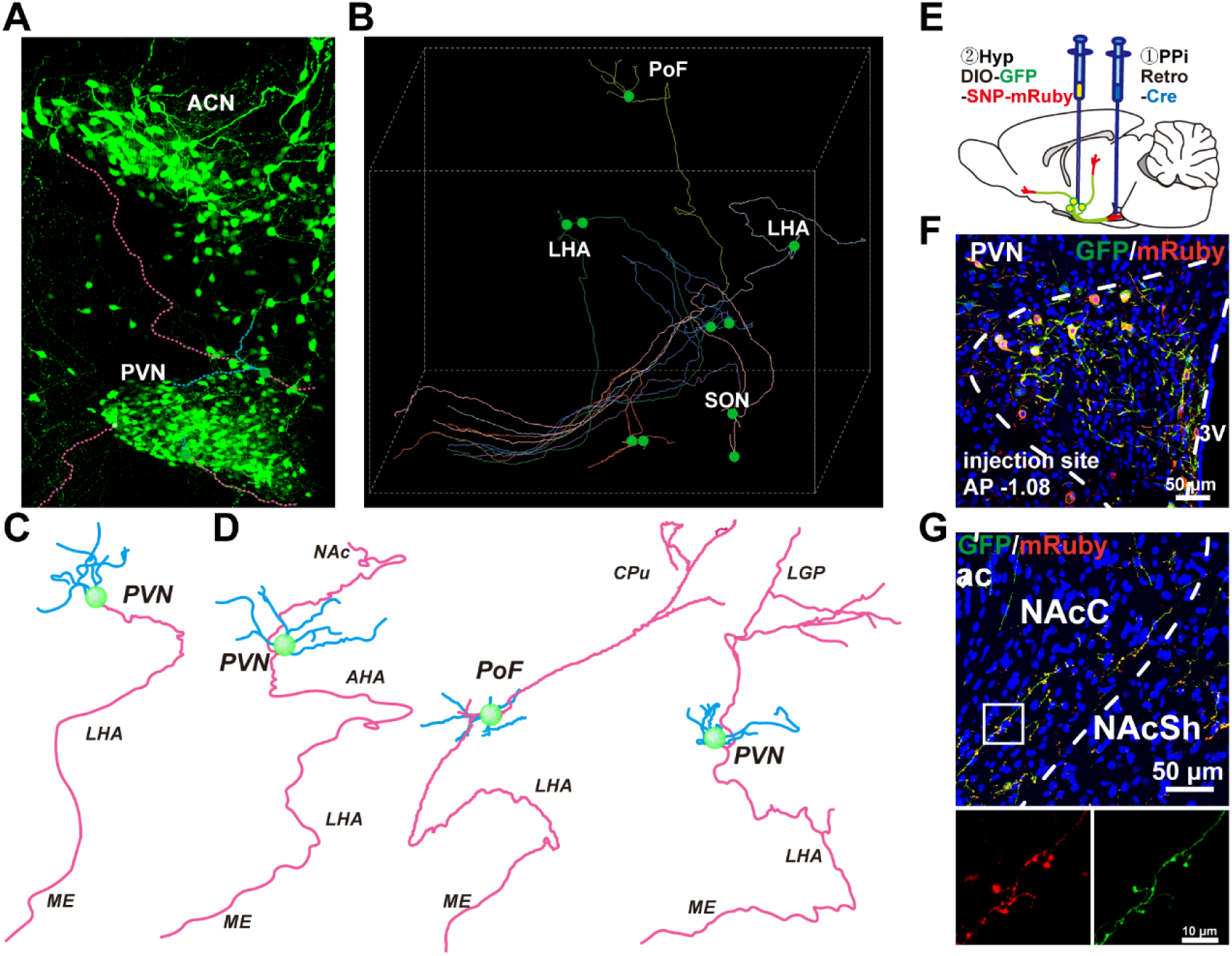
Collateral projections of MNCs in multiple extra-hypothalamus regions by single-cell fiber tracing and dual-labeling strategy. (A) Representative images of the single-cell fiber tracing in ACN and PVN. (B) Representative images showing the skeletonization of fiber tracing in regions containing PoF, LHA and SON. Green dots denote the somas; lines represent the fiber routes. (C and D) Representative images of single-cell tracing of the MNCs (46 cells) with annotation of fiber path. The green spheres represent the somas, while the magenta and blue lines represent the trajectory of axons and dendrites respectively. The collateral projections of PPi-projecting MNCs to the medium eminence (ME) or NAc, LGP and CPu were shown (D). Images were not in the same scale. (E) Collateral labeling of MNCs in PVN by dual-viral tracing strategy. (F) Labeling MNCs at the viral injection site in the PVN. (G) Distribution of GFP positive fibers and SNP-mRuby labelled axon terminals in NAc. Scale bar: 50 µm and 10 µm for high-magnification images of terminals.

### Magno-OXT Neurons Collaterally Projected to Multiple Brain Regions

OXT plays an important role in regulating diverse social behaviors, including pair bonding and affiliative behaviors (Burkett et al., 2016; Jurek and Neumann, 2018; Young and Wang, 2004). However, it remains controversial whether Magno-OXT or Parvo-OXT neurons are involved. To address this question, we generated an OXT-Cre rat line, in which a P2A-iCre cassette was inserted after the *OXT* gene using the CRISPR-Cas9 strategy and verified by Southern blot (Figure S5A-S5C). Next, injection of the Cre-dependent retrograde viral tracer (rAAV2-Retro-DIO-YFP) into the PPi, followed by OXT-neurophysin (OXT-NP) immunostaining, revealed that more than 95% of virally-traced cells in the PVN and SON were positive for OXT-NP, suggesting specific Cre expression in OXT neurons (Figure 5A, 5B and 5D). Conversely, approximately two thirds (64.6%) of OXT-NP positive cells were labelled by YFP in SON, suggesting a similar labeling efficiency observed in SD rats (Figure 5D). Moreover, OXT cells in the PMN and AMN were also successfully labeled (Figure S5D and S5E). By contrast, none of the Parvo-OXT cells in the PVN were labeled, verifying the specific labeling of Magno-OXT neuron ensemble by PPi-retrograde tracing (Figure 5C). Consistent with our above observations (Figures 3B and S4C), YFP^+^ fibers were also found in several brain areas, including the NAc, LS, Piri C, and CPu (Figure 5E). These data demonstrate that Magno-OXT neurons collaterally project to the extra-hypothalamic regions.

**Figure 5.**
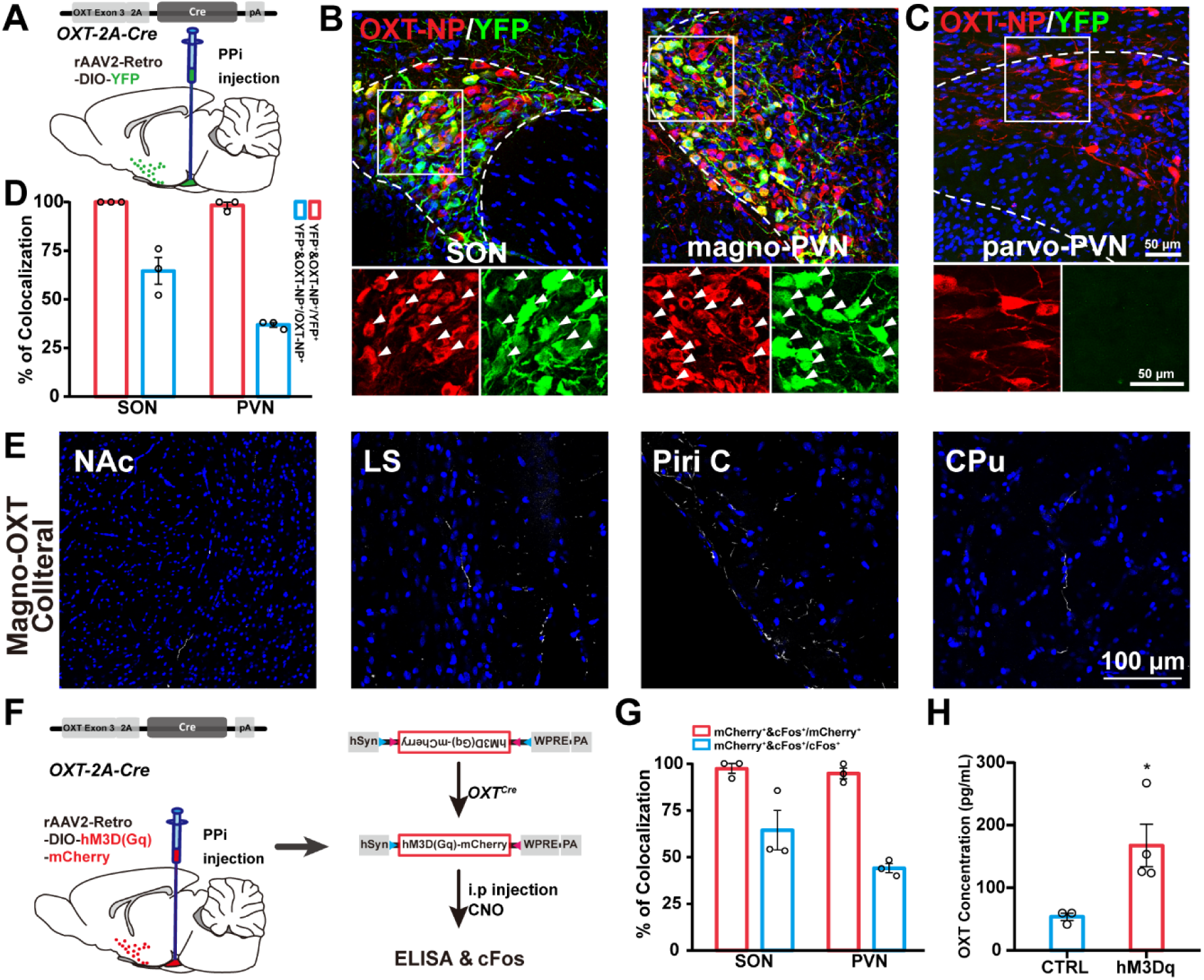
Chemogenetic activation of Magno-OXT neurons promotes oxytocin release. (A) Schematic injection and viral infection of rAAV2-Retro-DIO-YFP in OXT-Cre rats. (B) Representative images showing the validation of OXT-Cre line. Co-localization of YFP with OXT-neurophysin (NP) was indicated with white triangles. (C) The parvocellular neurons in the PVN were negative for YFP. (D) Histogram of the OXT-NP and YFP co-localization in the PVN and SON. (n=3 rats, 182 cells in the PVN, 257 cells in the SON). (E) Distribution of YFP-labelled fibers of magnocellular oxytocin (Magno-OXT) neurons in the extrahypothalamic regions including the NAc, LS, Piri C and CPu. (F) Diagram of chemogenetic activation of Magno-OXT neurons and the subsequent measurement. ELISA and cFos staining were carried out 90 min after the injection of CNO (1 mg/kg). (G) Histogram of the cFos and mCherry co-localization. 97.4% of the mCherry^+^ cells were positive for cFos. 64.5% of activated Magno-OXT neurons were mCherry^+^ in SON (n=3 rats, 335 cells) and 44.1% in PVN (n=3 rats, 344 cells). (H) Elevated oxytocin levels in the blood after CNO treatment as measured by ELISA (CTRL, 53.54±5.99 pg/mL, n=3 rats; Dq, 167.63±33.93 pg/mL, n=4 rats; p=0.0371, two-tailed *t*-test.). Scale bar: 50 μm.

### Selective Activation of Magno-OXT Neurons Enhanced Peripheral Oxytocin Release and Facilitated Central OXT-mediated Social Interactions

To selectively activate Magno-OXT neurons, we injected the rAAV2-Retro-DIO-hM3D (Gq)-mCherry (Retro-DIO-Dq), which expresses the engineered receptor exclusively activated by a designed drug clozapine-noxide (CNO) (Armbruster et al., 2007), into the PPi of OXT-Cre rats (Figure 5F). Consistent with data above (Figure 5B and 5C), the virus selectively labeled Magno-but not Parvo-OXT neurons. Immunostaining of c-Fos, an immediate neuronal activation marker, revealed that almost all retro-traced (mCherry^+^) cells (97.4±1.4%) in PMNs, AMNs and SMNs were positive for c-Fos after CNO administration (Figure 5G, S5F-S5H). Conversely, parvo-OXT neurons in the PVN were negative for c-Fos, suggesting that CNO selectively activated Magno-OXT neurons (Figure S5I). Next, to test the effects of activation of Magno-OXT neurons on peripheral OXT release, we measured the plasma OXT concentration and found that peripheral OXT levels were increased by three folds one hour after CNO treatment (Figure 5H). Therefore, the retrogradely transported chemogenetic tool provides an effective approach to selectively manipulate the Magno-OXT neuronal ensemble.

To further examine whether activation of Magno-OXT neurons alone induces central OXT related behaviors, in particular, social behaviors, male rats injected with Retro-DIO-Dq or control viruses were placed into an open-field (OF) apparatus to interact with a juvenile stimulus rat (Figure S6A). Compared to the controls, CNO treatment significantly increased the time of Retro-DIO-Dq infected rats socializing with the stimulus rats (Figure S6B-S6D). In a three-chamber apparatus, rats showed no preference to a certain chamber during the habituation and exploration stages (Figure 6A and 6C). However, at the social investigation stage, the time for the Dq group spent in the stimulus rat-containing chamber was significantly longer than the control group (Figure 6B and 6C). Moreover, Dq rats spent as twice time in the social zone (10 cm around the stimulus cage) as the controls and exhibited robustly elevated social index (duration in the social zone divided by the total duration in two zones) (Figure 6D and 6E), suggesting enhanced social interaction-associated chamber preference. In accordance with previous findings showing that OXT conveys social information through the olfaction (Oettl et al., 2016; Wesson, 2013), sniff counts were increased by two folds in Dq rats (Figure 6F). Notably, when the stimulus rat was replaced with a toy, Dq rats showed no increase of duration or sniff counts in the social zone upon CNO administration (Figure S6H-S6J), suggesting that OXT specifically facilitates social interactions. As peripheral OXT barely passes BBB (Leng and Ludwig, 2016; Mens et al., 1983), central OXT released from activated Magno-OXT neurons likely mediates these behavioral effects. Collectively, our data demonstrate that activation of Magno-OXT neurons induces both peripheral and central effects of OXT.

**Figure 6.**
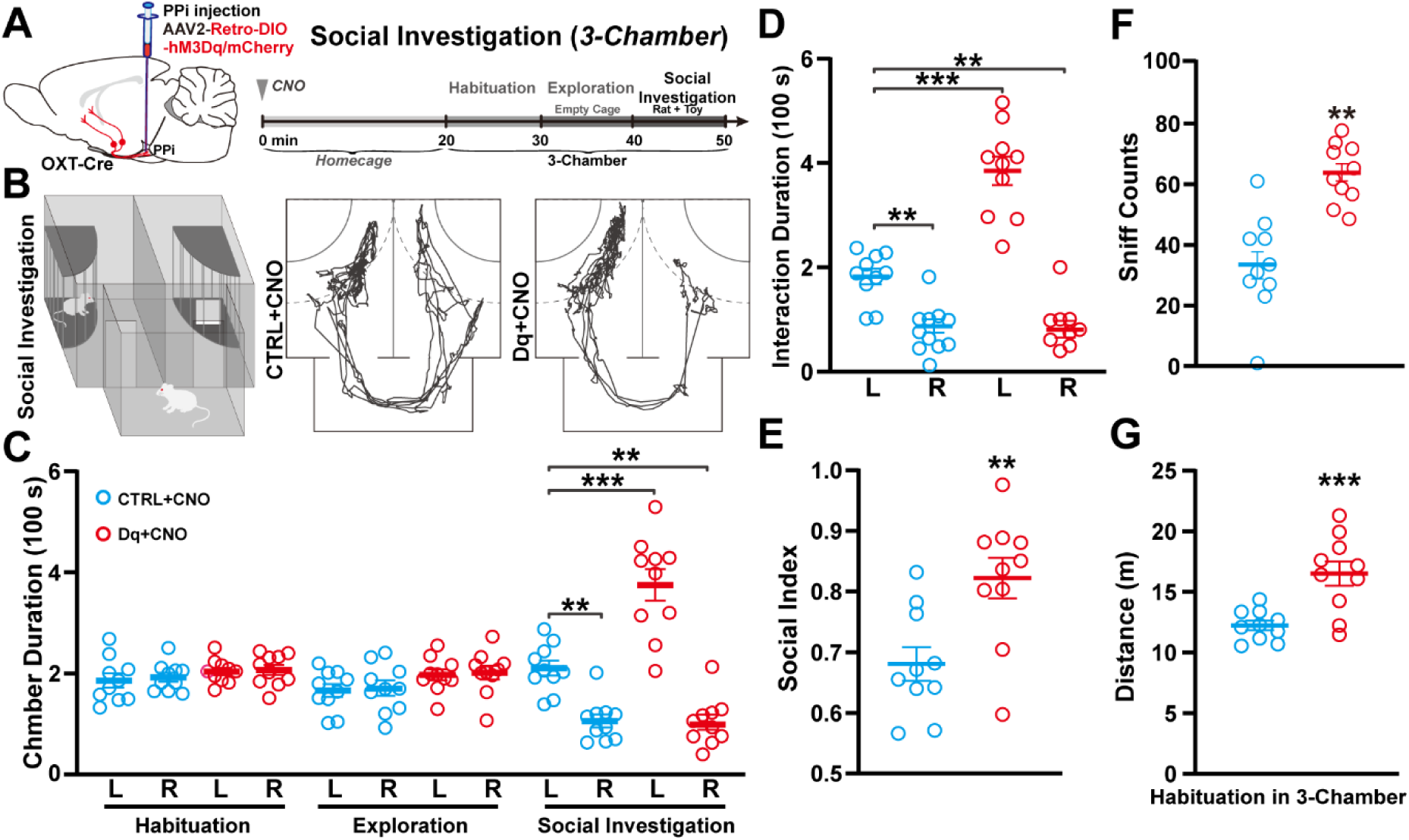
Chemogenetic activation of Magno-OXT neurons promoted social investigation. (A) Diagram of viral injection, drug administration and social investigation tests. Retro-DIO-hM3Dq viruses were injected into the PPi of OXT-Cre rats. The habituation, exploration and social investigation were proceeded 20 min after CNO administration. (B) Diagram and track plots showing the social test in a 3-chamber apparatus. (C) Duration of CTRL and Dq-injected rats in the left and right chambers at different stages. Activation of Magno-OXT neurons increased the duration in left chamber during the social investigation stage (CTRL, n=10 rats, social L-chamber 208.2 s; Dq, n=10 rats, social L-chamber 385.3 s). (D and E) Activation of Magno-OXT neurons increased the duration in the social zone. Social index was calculated as duration in the left social zone divided by the total duration in both zones (CTRL, n=10 rats, duration=176.1 s, index=0.45; Dq, n=10 rats, duration=371.9 s, index=0.79. Duration analysis, Fr(1,36)=112.2, p<0.0001; Fc(1,36)=27.48, p<0.0001, two-way ANOVA group × zone. Social index analysis, p=0.0044, two-tailed *t*-test.). (F) Increased sniff counts of Dq group after intraperitoneal administration of CNO. (CTRL, n=10 rats, 33.3; Dq, n=10 rats,67. p<0.0001, two-tailed *t*-test). (G) Increased total travel distance upon activation of Magno-OXT neurons during the habituation stage (CTRL, n=10 rats; Dq, n=10 rats. p=0.0009, two-tailed *t*-test.).

### Activation of Magno-OXT Neurons Promoted Locomotion via OXT Release in CPu

In measuring social behaviors, we noticed that CNO-treated Dq rats increased travel distances in the habituation stage, either in the OF or 3-chamber apparatus, suggesting that activation of Magno-OXT likely enhances locomotor activities (Figure 6G, S7E and S7K). Given that OXT neurons collaterally project to the CPu/LGP (Figure 5E), a region critical in motor regulation (Grillner et al., 2005), and OXTR is expressed in the CPu/LGP (Figure 7A), we hypothesized that striatum-projecting Magno-OXT neurons might be involved in the regulation of locomotion. As previous studies suggested that OXT may elicit dosage-dependent effects on locomotion (Klenerova et al., 2009; Uvnas-Moberg et al., 1994), we further tested the locomotion of the rats in a specific box (Figure 7B-7D). Measurement of the travel distance, line crossing and rearing behaviors revealed that CNO-but not saline-treated Dq animals showed elevated horizontal and vertical locomotion, whereas CNO-treated controls showed no changes (Figure 7E-7H). To examine whether the effects were modulated by OXTR mediated signaling, we infused L-368,899, a commonly used OXTR antagonist, into the CPu and observed that CNO-induced locomotion increase was abolished (Figure 7F-7H), suggesting a role of striatal OXT in the increased locomotion. To further test whether activation of Magno-OXT neuronal terminals induces similar effects, we directly infused CNO into the CPu to trigger release of OXT from local terminals. We did observe increased line crossing; however, the total travel distance and rearing were not significantly different (Figure 7I-7K), likely due to the sparsely distributed collaterals and/or insufficient terminal activation in the region. Together, these results demonstrate that activation of Magno-OXT neurons drives locomotor activity through striatal OXT signaling.

**Figure 7.**
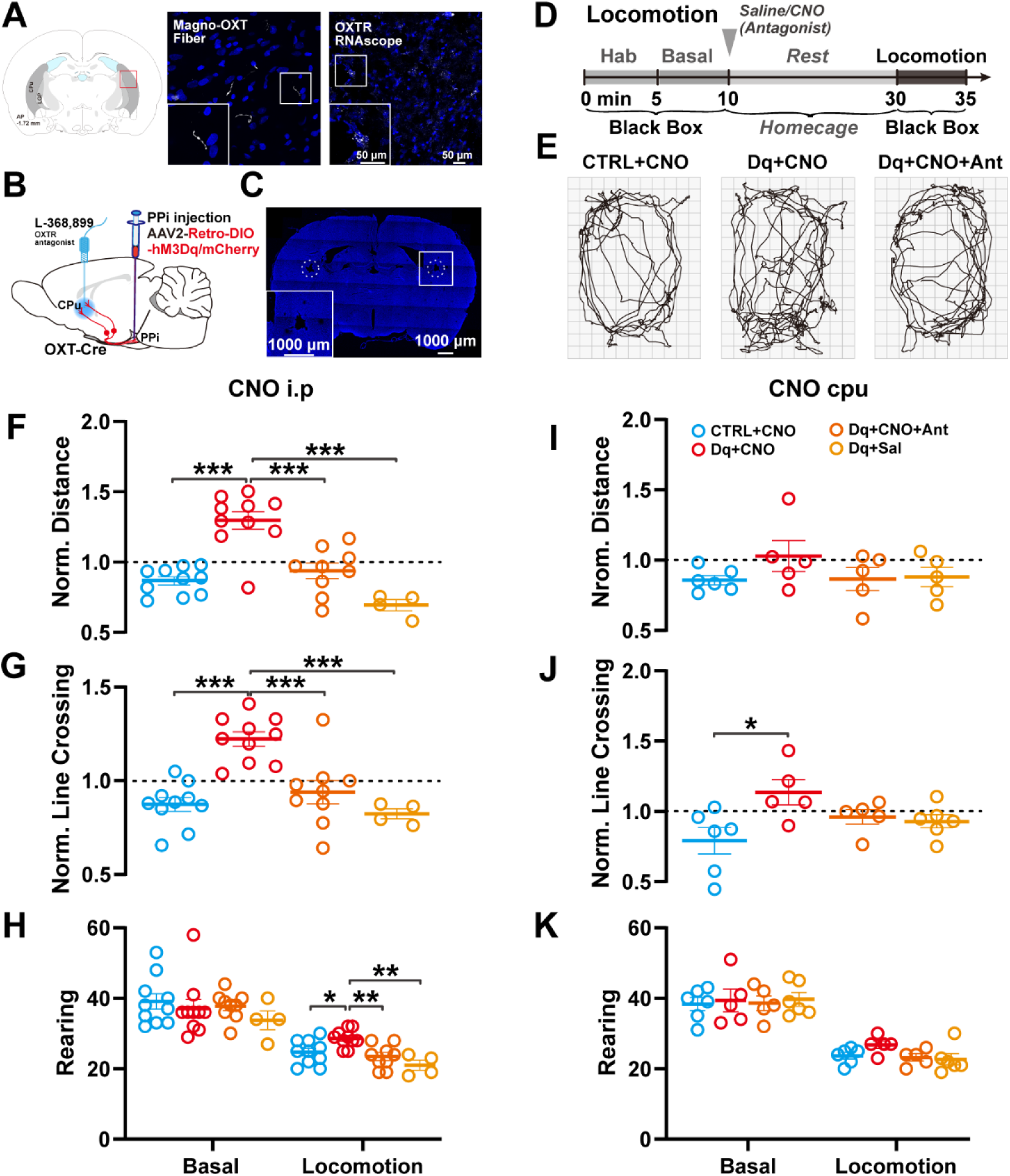
Chemogenetic activation of Magno-OXT neurons enhanced locomotion. (A) Left panel, a schematic of striatal slice; Middle panel, GFP^+^ fibers in the CPu and LGP of retrogradely-traced Magno-OXT neurons; Right panel, expression of OXT receptors in CPu by single molecule fluorescence *in situ* hybridization. (B-D) Diagram of drug administration and locomotion tests. Retro-DIO-hM3Dq was injected into the PPi of OXT-Cre rat, and microinfusion cannulas were implanted in bilateral CPu (B). The cannula tract was indicated with dotted line (C). Drugs were administrated following the habituation and basal stage. Locomotion was tested 20 min after administration of CNO/Saline in the presence or absence of OXTR antagonist L-368,899 (D). (E) Representative track plots of the locomotion activity of animals after different treatments. (F-H) Increased locomotion after CNO-induced activation of Magno-OXT neurons. Total travel distance (F), line-crossing (G) and rearing (H). OXTR antagonist abolished the locomotion-promoting effects (n=10, 10, 10 and 4 rats for CTRL, Dq+CNO, Dq+CNO+Antagonist and Dq+Saline groups, respectively. All data were normalized to the basal stage. Distance analysis, F=20.59, p<0.0001, one-way ANOVA. Line-crossing analysis, F=14.67, p<0.0001, one-way ANOVA.). (I-K) Effects of CNO-induced terminal activation of Magno-OXT neurons by infusion of CNO in CPu. Travel distance (I), line-crossing (J) and rearing (K) (n=6, 5, 5 and 5 rats for CTRL, Dq+CNO, Dq+CNO+Antagonist and Dq+Saline groups, respectively. Normalized to basal level. Distance analysis, F=1.133, p=0.3638, one-way ANOVA. Line-crossing analysis, F=3.663, p=0.3020, one-way ANOVA.). Scale bar: 50 μm (A) and 1000 μm (D).

## DISCUSSION

Using state-of-the-art techniques in viral tracing and brain-wide imaging, we systematically mapped the connectivity of the HNS and uncovered the anatomical links between multiple hypothalamic MNC nuclei and the neurohypophysis. These data lay the groundwork for fully understanding the architecture and regulation of the neuroendocrine networks in the future. We further reveal that subsets of MNCs collaterally project to both neurohypophysis and multiple brain areas, providing structural evidence for peripheral and central release of neuropeptides from MNCs. In addition, we demonstrate that activation of Magno-OXT neurons alone not only elevates peripheral OXT levels but also promotes central OXT release to facilitate social interactions and locomotion. Together, these findings suggest that MNCs play important roles in maintaining homeostasis and regulating behavioral responses, and such actions were achieved through coordinated peripheral and central release of peptides.

Earlier attempts to map the HNS with retrograde tracers provided important information on the general structures of the HNS (Fisher et al., 1979; Kelly and Swanson, 1980; Peterson, 1966). However, limited labeling efficiency and imaging resolution prevented a clear visualization of the system (Fisher et al., 1979; Y et al., 1988). Retrograde viral tracing from the PPi is an ideal approach to label MNCs, together with advanced imaging techniques, we fully reconstructed the architecture of the HNS with high resolution for the first time. The reconstructed 3D atlas uncovers the complexity of the HNS and expands the composition of the HNS, which has not been recognized previously. It should be noted that the predominantly-studied PVN and SON only harbor approximately a half of the MNCs and functions of the rest of MNCs remain largely unknown. It would be interesting to investigate whether MNCs in AMN and SMN play complementary or opposite roles to those in PMN. On the other hand, the reconstructed atlas is unable to specifically delineate the distribution and projections of Magno-AVP or Magno-OXT neurons. Given the functional importance and segregation of AVP and OXT in the HNS (Heinrichs et al., 2009; Meyer-Lindenberg et al., 2011; Neumann and Landgraf, 2012; Stoop, 2012), it would be important to selectively map the connectomes of Magno-OXT and Magno-AVP neurons in the brain.

MNCs have been regarded as the output or “motor” neurons that send unipolar axons and secret neuropeptides into the bloodstream (Buijs and Van Eden, 2000; Harris, 1955a; Swanson, 2012; Watts, 2005). Using single-cell fiber tracing, we found that a fraction of MNCs collaterally projects to both the neurohypophysis and the extrahypothalamic regions. These data suggest that in addition to maintain homeostasis by peripheral hormone secretion, MNCs also coordinate related behaviors by central peptide release. For example, the full engagement of Magno-OXT neurons during parturition and lactation, which reaches maximal OXT release into the bloodstream, may also promote central release through the collaterals to enhance maternal behaviors and mother-infant affiliation (Kendrick, 2000; Young and Wang, 2004). To support this notion, a study using dual-retrograde labeling observed that OXT neurons in the PVN collaterally project to the NAc in prairie voles, likely promoting pair bonding effects (Ross et al., 2009). A recent study confirming the modulatory role of Magno-OXT also showed that Magno-OXT neurons in the SON collaterally project to the septum to attenuate conditioned fear (Menon et al., 2018). Meanwhile, a limitation of the current study is that single-cell fiber tracing was carried out in a small number of MNCs due to the high density of traced cells. Future studies will use the newly-developed dual-AAV suites to sparsely label MNCs in different nuclei to further map their projections in full spectrum (Lin et al., 2018).

The wide distributions of MNCs in a variety of hypothalamic nuclei raised a challenge in labeling and manipulating the neuronal ensemble. Knobloch et al. carried out multiple injections to label the OXT neurons and uncovered extensive projections of OXT neurons to the brain (Knobloch et al., 2012). However, these results may have obscured the contribution of Parvo-OXT neurons, as both Magno-OXT and Parvo-OXT neurons can be infected. Using rabies viruses tracing, the authors observed collateral projections of Magno-OXT neurons to the amygdala (Knobloch et al., 2012). We further clarified that Magno-OXT neurons collaterally project to multiple extrahypothalamic regions, including the Piri C, Audi C and NAc (Figure 5E). These findings may help to explain the roles of central OXT in regulating social recognition, learning and memory as well as social rewards (Choe et al., 2015; Marlin et al., 2015). Intriguingly, although OXT has been shown to gate social rewards at both NAc and VTA (Dolen et al., 2013; Hung et al., 2017), we found that MNCs and/or Magno-OXT neurons specifically project to NAc but not VTA. This is consistent with a recent study demonstrating that Parvo-OXT neurons innervate VTA and regulate the activity of VTA neurons in mice (Xiao et al., 2017).

We further demonstrate that retrograde tracing from the PPi followed by chemogenetic stimulation is an efficient approach to selectively manipulate the activities of Magno-OXT neurons without affecting Parvo-OXT neurons. Activation of Magno-OXT neurons alone promotes peripheral release of OXT and enhances central OXT-mediated social interactions and locomotion simultaneously. While the vast majority of studies have mainly focused on OXT neurons (including both Magno-OXT and Parvo-OXT neurons) in the PVN (Choe et al., 2015; Marlin et al., 2015; Oettl et al., 2016; Xiao et al., 2017), this study made a first attempt to clarify functions of individual OXT neuronal subgroups and specifically demonstrated that Magno-OXT neurons contribute to the central actions of OXT. Although injection of CNO at CPu only led to moderately increased locomotion, local infusion of OXTR antagonist significantly attenuated the locomotion increase. Together with previous observation that *OXT* knockout mice showed decreased tendency in travel distances in a social approach test (Crawley et al., 2007), these findings suggest that OXT in CPu enhances locomotion. We postulated that that increased locomotion may contribute to active exploration to facilitate social investigation (Blume, Eur J Neurosci, 2008). It is possible that the activation of Magno-OXT neuron elicits central OXT release, which drives the animals to be more active and exploratory to facilitate social interactions (Calcagnoli et al., 2014; Oettl et al., 2016).

HNS serves as the Rosetta stone in the regulation of neuroendocrine networks. Our work provides important references for further illumination of the hard-wiring circuits and functions of MNCs. The extensive collaterals of Magno-OXT neurons and the effects elicited upon their activations suggest that they directly modulate neural circuits related to social behaviors. The current work provides new insights and clues to better understand the mechanisms underlying a wide range of mental disorders, such as depression, autism and anxiety, in which dysfunctions of the oxytocinergic transmission have been implicated.

## AUTHOR CONTRIBUTIONS

Conceptualization, Z. G., B. Z., S. D.; Methodology, B. Z., L. Q., H. N.; Software, A. L., H. G.; Validation, W. X., L. Q.; Formal Analysis, B. Z., L. Q., W. X., H. N.; L. C., F. W., W. M.; Investigation, B. Z., L. Q., F. W.; Resources, A. L., H. G.; Data Curation, B. Z., A. L.; Z. G., S. D., Writing – Original Draft, Z. G., B. Z., L. Q., A. L.; H. G.; Writing – Review & Editing, Z. G., L. Q., W. X.; Visualization, B. Z., W. X.; Supervision, Z. G., S. D., H. G.; Project Administration, Z. G., B. Z., L. Q., W. X.; Funding Acquisition, Z. G., S. D., H. G., A. L. and B. Z..

## ACKNOWLEDGMENTS

We thank Dr. Harold Gainer at National Institute of Health for generously providing the neurophysin antibodies, Dr. Gareth Leng for valuable suggestions on manuscript revision, Dr. D. F. Swaab and Aimin Bao for valuable suggestions on experimental design, Dr. Xing Zhang Hangjun Wu for technical assistance with the image processing. This work was supported by the National Key Research and Development Program of China (2016YFA0501000), National Natural Science Foundation of China (31671057, 61721092, 81821091, 31490592, 81527901, 91832000, 91732304), Science and Technology Planning Project of Guangdong Province (2018B030331001), and the Fundamental Research Funds for the Central Universities (2019FZA7009).

## MATERIALS AND METHODS

### KEY RESOURCES TABLE

| REAGENT or RESOURCE | SOURCE | IDENTIFIER |
| --- | --- | --- |
| **Antibodies** |  |  |
| Normal goat serum | Jackson ImmunoResearch | 005-000-121 |
| Normal donkey serum | Jackson ImmunoResearch | 017-000-121 |
| Rabbit anti-GFP (1:2000) | Abcam | Ab6556 |
| Goat anti-GFP (1:1000) | Abcam | Ab5450 |
| Rabbit anti-Arginine Vasopressin (1:20000) | Phoenix Biotech | H-065-07 |
| Rabbit anti-Oxytocin (1:10000) | Phoenix Biotech | H-051-01 |
| Mouse anti-Vasopressin-associated Neurophysin (1:100) | Gift from Dr. Harold Gainer | PS41 |
| Mouse anti Oxytocin-associated Neurophysin (1:200) | Gift from Dr. Harold Gainer | PS38 |
| Rabbit anti-Fluorescent Gold (1:500) | Merck Millipore | AB153-I |
| **Bacterial and Virus Strains** |  |  |
| rAAV2/2-Retro-CMV_bGI-EGFP-WPRE-pA | Taitool | S0263 |
| rAAV2/2-Retro-hSyn-Cre-WPRE-pA | Taitool | S0278 |
| AAV2/9-hSyn-DIO-mGFP-2A-Synaptophysin-mRuby | Taitool | S0250 |
| rAAV2/2-Retro-hSyn-DIO-hM3D(Gq)-mCherry-WPRE-pA | Taitool | S0192 |
| rAAV2/2-Retro-hSyn-DIO-mCherry-WPRE-pA | Taitool | S0197 |
| **Chemicals, Peptides, and Recombinant Proteins** |  |  |
| Cholera toxin subunit B (recombinant), Alexa Fluor® 488 conjugate (CTB-488) | Life Technologies | C22841 |
| Prolong® Gold antifade reagent | Life Technologies | P10144 |
| Fluoro-Gold™ | FluoroChrome | 223769-64-0 |
| Clozapine N-oxide | Sigma-Aldrich | C0832 |
| **Critical Commercial Assays** |  |  |
| Oxytocin ELISA kit | Enzo | ADI-901-153A |
| 200 mg C18 columns | Sep-Pak | 043038348A |
| **Experimental Models: Organisms/Strains** |  |  |
| Oxytocin-Cre (OXT-Cre) rats | Biocytogen (Nanjing, China) | EGE-ZY-028 |
| **Software and Algorithms** |  |  |
| ImageJ | NIH | https://imagej.nih.gov/ij/ |
| Amira | ThermoFisher Scientific | https://www.thermofisher.com/cn/zh/home/industrial/electron-microscopy/electron-microscopy-instruments-workflow-solutions/3d-visualization-analysis-software/amira-life-sciences-biomedical |
| Imaris | OXFORD Instruments | https://imaris.oxinst.com/ |
| SigmaPlot | SYSTAT | https://systatsoftware.com/products/sigmaplot/ |
| Prism | GraphPad | https://www.graphpad.com/scientific-software/prism/ |

## LEAD CONTACT AND MATERIALS AVAILABILITY

### Materials Availability Statement

Further information and requests for resources and reagents should be directed to and will be fulfilled by the Lead Contact, Zhihua Gao.

## EXPERIMENTAL MODEL AND SUBJECT DETAILS

### Animals

All animal experiments were approved by Zhejiang University Animal Facilities and followed the guidelines. Oxytocin-Cre (OXT-Cre) rats were generated by Biocytogen (Nanjing). Wild type Sprague Dawley (SD) rats were maintained in the Lab Animal Center at Zhejiang University and group housed on a 12-hour light cycle (lights off at 7:00 PM). Behavioral tests were performed with rats aged 15-20 weeks during the dark cycle.

## METHOD DETAILS

### Stereotaxic injection

Rats aged 12-16 weeks (260-280 g) were used for injection. The rats were anesthetized with 1% pentobarbital and placed on a heating pad to maintain the body temperature. Viruses were injected using a 2.5 µl Hamilton syringe driven by a pressure microinjector (KD Scientific) and guided with a stereotaxic instrument (David Kopf Instruments). The needle was inserted to the target locations based on the stereotaxic atlas (Paxinos and Watson, 2006; Swanson, 2004), through a drilled hole in the brain skull. Virus was infused at a slow rate over 10 min. To allow diffusion and reduce backflow, the syringe was maintained at the target position for 10 min at the end of injection, then uplifted 50 µm and maintained for another 3 min before complete withdrawal.

For retrograde tracing, 1 µl rAAV2/2-Retro-CMV_bGI-EGFP was injected into the posterior lobe of the pituitary (PPi) (AP, −5.80 mm; ML, +0.05 mm; DV, −10.20 mm) at 100 nl/min. The rats were sacrificed or further processed after 2 weeks.

For collateral projection analysis, 1 µl rAAV2/2-Retro-hSyn-Cre was injected into the PPi at 100 nl/min, followed by 200 nl AAV2/9-hSyn-DIO-mGFP-2A-Synaptophysin-mRuby injection 1 week later into the PVN (AP, −0.72 mm; ML, +0.60 mm; DV, −7.20 mm) at 20 nl/min. The virus was allowed to express for another 4 weeks.

To label neuroendocrine cells, rats injected with rAAV2-Retro-GFP received an i.p. injection of 4% Fluoro-Gold in 0.9% NaCl (30 mg per kg body weight) and sacrificed 10 days later.

To specifically retrogradely trace the magnocellular OXT neurons, 1 µl rAAV2/2-Retro-hSyn-DIO-YFP was injected into the PPi of OXT-Cre rat at 100 nl/min. The virus was allowed to express for another 2 weeks.

For chemogenetic activation, 1 µl rAAV2/2-Retro-hSyn-DIO-hM3D(Gq)-mCherry-WPRE-pA or control virus rAAV2/2-Retro-hSyn-DIO-mCherry-WPRE-pA was injected into the (PPi) at 100 nl/min. 2 weeks post injection, cannulas were implanted bilaterally at CPu (AP, −1.80 mm; ML, ±3.60 mm; DV, −5.40 mm). The rats were allowed to recover for 1 week before handling.

### Behavioral tests

All behavioral tests were performed within a noise cancellation room maintained at 25 ℃. The rats were fed in the same room for 7 days to acclimate to the environment after surgery. For all experiments, experimenters were blinded to genotype or experimental manipulation. All the apparatuses and cages were sequentially wiped with 70% ethanol and ddH_2_O then air-dried between stages.

At the end of behavioral tests, 1 mL blood was collected from the left heart ventricle of deeply anesthetized rats for ELISA. After blood collection, rats were perfused with 4% paraformaldehyde followed by *post-hoc* analysis to confirm the injection sites and cannula location.

Clozapine N-oxide (CNO) (in 500 μL 0.9% saline) was intraperitoneally injected 20 min before behavioral test at 1 mg per kg body weight. For CPu microinfusion, the infusion catheter was designed to be 200 μm longer than the guiding cannula and connected to a 2.5 μL Hamilton syringe via a polyethylene tube. 1000 nL ACSF or drugs were infused into bilateral CPu (500 nL each side) with a manual microinfusion pump (RWD, 68606) over 3 min. The catheter was kept in the CPu for another 1 min to maintain complete diffusion. The dosage we used was listed as below. CNO was diluted in ACSF to make a 5 μM working solution. L-368,899 was dissolved in ACSF to make a 2 μg/mL working solution.

Movements were recorded using a Sony camera connected to the Any-Maze system. The duration spent exploring different areas was automatically acquired by Any-Maze. The sniffing was defined as valid when the noses of subject and object rats touched or oriented towards each other within 2 cm, the counts were manually calculated by two experimenters unaware of the treatments. The rearing was defined as the rat standing with the hindlimbs and also manually calculated.

#### Locomotion test

A black box of 33 cm in width and 45 cm in length was used to assess the locomotion activity of rats. A grid consisting of 3 × 3 cm square was applied to trace the line-crossings and distance during movement. The subject rat was first allowed to habituate in the box for 5 min and the basal locomotion level of the animal was measured for 5 min right after the habituation. The animal was then put back to the homecage, where drugs were given via intraperitoneal injection or CPu microinfusion. 20 minutes after the injection, the subject rat was placed back to the black box and locomotor activity was measured for 5 min. The total travel distance and line-crossings were calculated by Any-Maze to indicate the horizontal locomotion. The rearing was manually counted to indicate the vertical locomotion.

#### Open field social test

The open field apparatus was 100 × 100 × 40 cm (width × length × height).

The subject rat was exposed to the empty apparatus for 10 min. Then a transparent Plexiglas cylinder of 20 cm in diameter was placed at one corner of the apparatus. The rat was allowed to explore for 10 min to acclimate to the environment. Lastly, a virgin male SD rat was introduced into the cylinder cage and the vicinity (10 cm) of the cage was defined as social zone. The subject rat was allowed to explore for another 10 min. The total travel distance and duration in the social zone were calculated by Any-Maze.

#### 3-chamber social test

The T-shaped three-chamber apparatus was composed of two 27 × 40 × 40 cm (width × length × height) side chambers and one 15 × 40 × 40 cm (width × length × height) shuttle chamber connecting side ones. Two identical cages 15 cm in diameter were used as stimulus cages and the vicinity (10 cm) of the cages was defined as social zone.

For the first stage (habituation), the subject rat was placed in the shuttle chamber with head orienting the right chamber and allowed to explore the apparatus freely for 5 min to acclimate. Then, for the second stage (exploration), two stimulus cages were placed at the corner of each side chamber. The rat was allowed to explore the apparatus containing empty cages for 10 min. For the third stage (novelty), we introduced a toy into the cage of the right chamber while left the other cage empty to test the effect on novel object exploration. The subject rat was allowed to freely explore the apparatus for another 10 min.

A parallel paradigm was adopted from the above to test the effect on social investigation. The habituation and exploration stages were repeated. At the third stage (sociability), a virgin male SD rat was introduced into the stimulus cage in the left chamber and a different toy in the other cage. The total travel distance and duration in the social zone were calculated by Any-Maze. The sniff counts were manually calculated based on the video. The social index is calculated as DurationL/(DurationL+DurationR), an index of the preference to the left social zone.

### ELISA

Blood was temporarily stored in iced tubes pre-coated with heparin, protease inhibitor cocktail and EDTA. After immediate centrifugation at 2000 g at 4 ℃ for 2 min, the supernatants (serum) were transferred, aliquoted and stored at −80 °C until use.

Serum oxytocin levels were measured using a commercially available oxytocin ELISA kit combined with C18 column extraction to avoid interference from other components. Briefly, the 200 mg C18 columns were equilibrated with 1 mL acetonitrile, followed by 20 mL 0.1% trifluoroacetic acid (TFA). 200 μL serum was mixed with an equal volume of 0.1% TFA and centrifuged at 16000 g at 4 ℃ for 15 min. The supernatants were applied to the column and washed with 20 mL 0.1% TFA. Samples were eluted from the column using 1 mL 95% acetonitrile/5% of 0.1% TFA and evaporated with nitrogen.

Samples were reconstituted with 250 μL Assay Buffer from the ELISA kit and analyzed according to the product manual. Briefly, the oxytocin standard samples were prepared via gradient dilution and corresponding diluents were loaded into appropriate wells. Testing samples were also added to different wells followed by Conjugate and Antibody incubation at 4 ℃ overnight. After washing in Wash Solution 3 times, pNpp Substrates were added to the well and incubated at room temperature for 1 h, followed by the addition of Stop Solution. The plate was immediately read at 405 nm (correction between 570 nm and 590 nm) and the oxytocin content was calculated based on the standard curve and the optical density of the samples.

### Histology and imaging

#### Immunofluorescence analysis

Animals were perfused with normal saline and 4% paraformaldehyde (PFA) sequentially. The dissected brains and pituitary glands were post-fixed in 4% PFA, followed by dehydration in 30% sucrose. Antigen retrieval of brain slices was carried out as described previously (Gao et al., 2012). The brain slices were permeabilized in 0.05% Triton X-100 in Tris-buffered saline, blocked with 100 mM glycine and 5% bovine serum albumin (BSA) containing 5% normal donkey serum. Tissue sections were subsequently incubated with diluted primary and secondary antibodies as indicated, nuclei stained with 6-diamidino-2-phenylindole (DAPI), and slides mounted with antifade reagents. Images were acquired using the Olympus FV1200 confocal microscope.

### 3-D reconstruction with fMOST (fluorescence micro-optical sectioning tomography)

#### Tissue preparation for fMOST

All histological procedures were performed as described previously (Gong et al., 2016; Yang et al., 2013). Rats were anesthetized and perfused with 0.01 M PBS (Sigma-Aldrich Inc., St. Louis, US) and PBS containing 4% paraformaldehyde (PFA). Brains were carefully dissected and post-fixed in 4% PFA for 24 h. Fixed brains (n=5) were transferred to 0.01 M PBS and incubated at 4 ℃ overnight and dehydrated in graded ethanol solutions. Then, individual brain was impregnated with Glycol Methacrylate (GMA, Ted Pella Inc., Redding, CA) and embedded in a vacuum oven.

#### Whole-brain imaging

The whole-brain imaging process was based on a homemade Brain-wide Position System (BPS) (Gong et al., 2016), with a simultaneous propidium iodide (PI) staining to label the cytoarchitectonic landmarks and a brain-wide fluorescence micro-optical sectioning tomography (fMOST) via structured illumination. For whole-brain imaging, each sample was immobilized in a water bath on a three-dimensional nano-precision translation stage. The brain sample was sectioned coronally at an interval of 2 µm in an antero-posterior (AP) direction to achieve the axial scan, counterstained in PI, followed by fluorescence image acquisition via mosaic scan. The combination of mosaic x-y-scan and axial z-scan produced continuous dataset at a voxel resolution of 0.32 × 0.32 × 2 μm^3^, which allowed 3D reconstruction. For individual rat brain, approximately 8000 coronal slices were sectioned from an intact adult rat brain, with the original data size of 30 TB.

#### Image pre-processing

The original images were collected and saved in TIFF formats with high fidelity. Image preprocessing including mosaic stitching and illumination correction were performed on dual-channel images (PI and GFP). Briefly, the mosaics of each coronal section were stitched to obtain entire section images. Lateral and axial corrections were performed by calculating mean intensity along each direction and quantifying the average grey-scale values, respectively. Image preprocessing was implemented in C++, parallelly-optimized with the Intel MPI Library (v3.2.2.006). Images were executed on a computing server and saved at 8-bit depth in LZW-compression TIF format or transformed to TData format (Li et al., 2017).

#### Registration and reconstruction of the hypothalamo-neurohypophysis system

The reconstructions of all the projection neurons were completed interactively in Amira 3D/4D^+^ visualization and analysis software (Thermo Fisher Scientific, USA). 10 × 10 × 32 μm^3^ resampled datasets were loaded into the Amira segmentation module for surface generation. For the absence of 3D high-definition SD rat brain atlas, we adopted Swanson’s atlas to register the brain regions or nuclei. And the boundaries of magnocellular aggregates were defined according to Swanson’s atlas and PI staining cytoarchitectonic landmarks.

#### Cell-counting and soma volume analysis

Each GFP-positive region was extracted into single file from raw 3D datasets for further analysis. GFP-positive cells were semi-automatically counted and manually verified in Imaris. The percentage of GFP^+^ cells within individual nucleus/region was calculated and normalized to the total number of GFP^+^ cells. The soma volume was automatically acquired with Imaris based on the same parameters. The anatomical boundaries and hallmarks were identified based on the Brain Maps III: Structure of the Rat Brain (Swanson, 2004).

#### Long-distance neuron reconstruction

A TData plugin was used to convert the high-resolution, whole-brain datasets into individual data blocks for subsequent processing in Amira (Li et al., 2017; Lin et al., 2018). The dendrites and/or axons in each sub-volume were traced interactively using the Filament Editor module. The contrast and thickness were adjusted to extract the fluorescent signals from the background. The same parameters were set to data blocks of one sample to maintain the reproducibility. Segments with high signal-noise ratio (SNR) were traced automatically on a 2D plane. Semi-automatic tracing strategy was applied to those with low SNR. The initial nodes of fibers were determined by experienced annotators and Amira semi-automatically calculated the path between nodes. The orientation consistency, fluorescence intensity and structural features were taken into consideration to determine the connection between related nodes missing a certain segment. All the results were further checked by an independent annotator.

#### Fiber density calculation

Fiber density calculation was performed for specific brain regions including the auditory cortex, BLA, CEA, CPu, LGP, LS, motor cortex, NAc, piriform cortex and VTA, with left and right halves calculated separately. The entire images were resampled to the resolution of 0.96 × 0.96 × 2 μm^3^ and imported to Amira for segmentation. The voxels with gray value larger than 15 were detected as GFP signals. Fiber density was calculated as the percentage of signal voxels in segment voxels.

## QUANTIFICATION AND STATISTICAL ANALYSIS

All the measurements were exported to the GraphPad Prism 8 or SigmaPlot 12 and analyzed by *t*-test, one-way ANOVA or two-way ANOVA according to the forms of the data. Nonparametric tests were used if the data did not match assumed Gaussian distribution. Data were presented as Mean ± SEM, with statistical significance taken as *p < 0.05, **p < 0.01, and ***p < 0.001.

The statistical details of experiments were attached to corresponding legends. Briefly, for histology analysis, *t*-test was used, n value was from n=3 to n=5 (rats) or from n=30 to n=100 (cells). For behavioral test analysis, *t*-test, one-way ANOVA and two-way ANOVA were used, as appropriate. The group size ranged from n=4 to n=10.

