## Supplemental Figures for "Reconstruction of the Hypothalamo-Neurohypophysial System and Functional Dissection of Magnocellular Oxytocin Neurons in the Brain"

### SUPPLEMENTAL INFORMATION

### SUPPLEMENTAL FIGURES

#### Supplemental Figures and Legends

**Figure S1**, related to Figure 1. The distribution of GFP-labelled cells in the hypothalamus at the macroscopic scale

**Figure S2**, related to Figure 1. Retrogradely labelled magnocellular neurons were neuroendocrine cells

**Figure S3**, related to Figure 1. AVP and OXT heterogeneity in different magnocellular nuclei

**Figure S4**, related to Figure 2, 3 and 4. Analysis of MNCs morphology in 3D and single cell fiber tracing in the MNs

**Figure S5**, related to Figure 5. Generation and validation of OXT-2A-Cre rats and chemogenetic virus

**Figure S6**, related to Figure 6 and 7. Novel object investigation was not affected by chemogenetic activation of Magno-OXT neurons

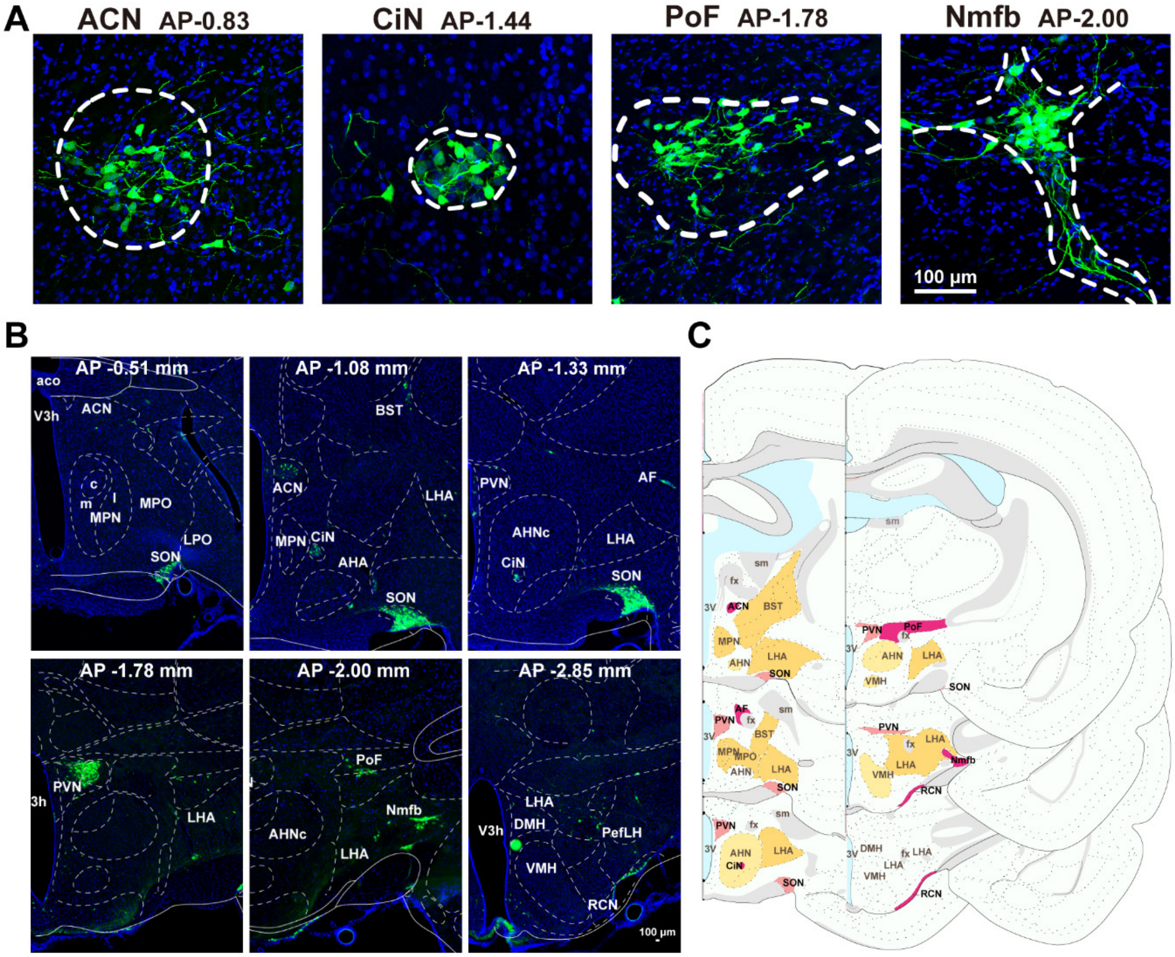
A
ACN AP-0.83
CiN AP-1.44
PoF AP-1.78
Nmfb AP-2.00
100 µm
B
AP -0.51 mm
AP -1.08 mm
AP -1.33 mm
AP -1.78 mm
AP -2.00 mm
AP -2.85 mm
C

**Figure S1**

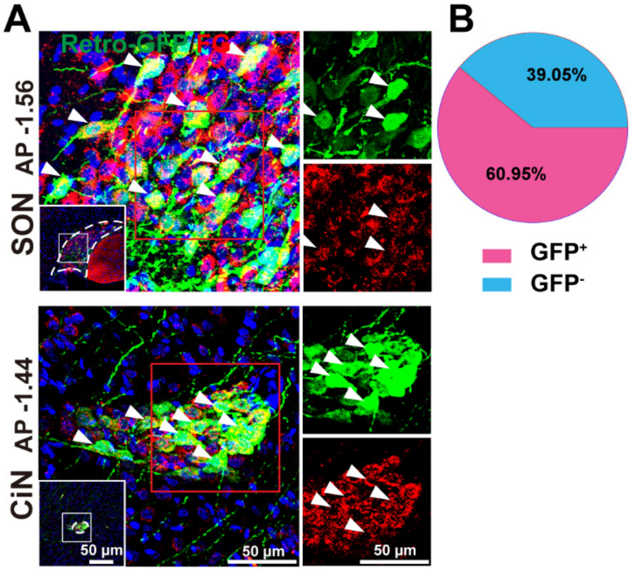
A
B
Retro-GFP/FG
SON AP -1.56
CiN AP -1.44
50 μm
50 μm
50 μm
39.05%
60.95%
GFP+
GFP-

**Figure S2**

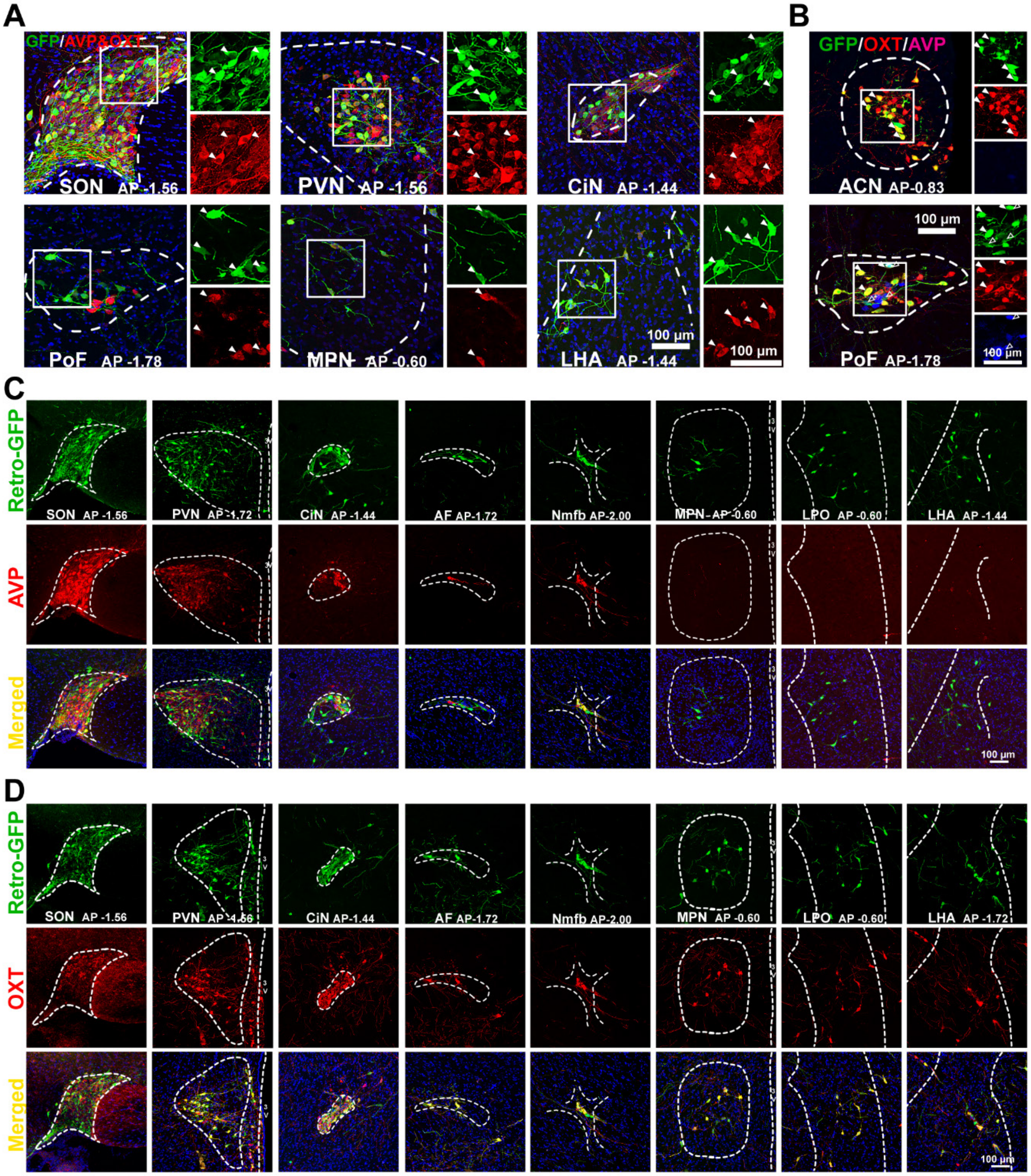
A
GFP/AVP&OXT
SON AP -1.56
PVN AP -1.56
CiN AP -1.44
PoF AP -1.78
MPN AP -0.60
LHA AP -1.44
100 µm
100 µm
B
GFP/OXT/AVP
ACN AP-0.83
PoF AP-1.78
100 µm
100 µm
C
Retro-GFP
AVP
Merged
SON AP -1.56
PVN AP -1.72
CiN AP -1.44
AF AP-1.72
Nmfb AP-2.00
MPN AP -0.60
LPO AP -0.60
LHA AP -1.44
100 µm
D
Retro-GFP
OXT
Merged
SON AP -1.56
CiN AP-1.44
AF AP-1.72
Nmfb AP-2.00
MPN AP -0.60
LPO AP -0.60
LHA AP -1.72
100 µm

**Figure S3**

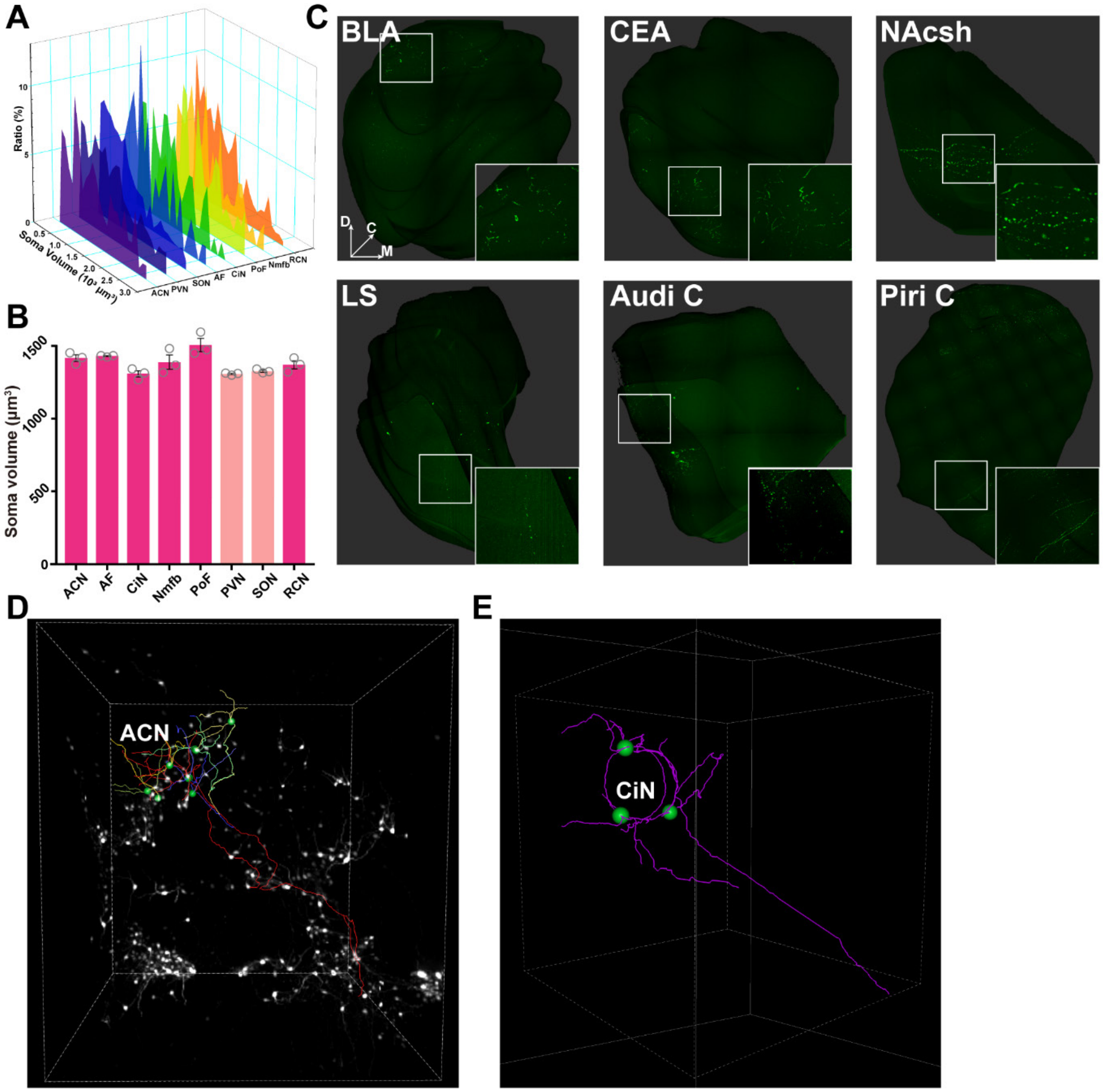
A
Ratio (%)
Soma Volume (10³ μm³)
ACN
PVN
SON
AF
CiN
PoF
Nmfb
RCN
B
Soma volume (μm³)
C
BLA
CEA
NAcsh
LS
Audi C
Piri C
D
C
M
D
ACN
E
CiN

**Figure S4**

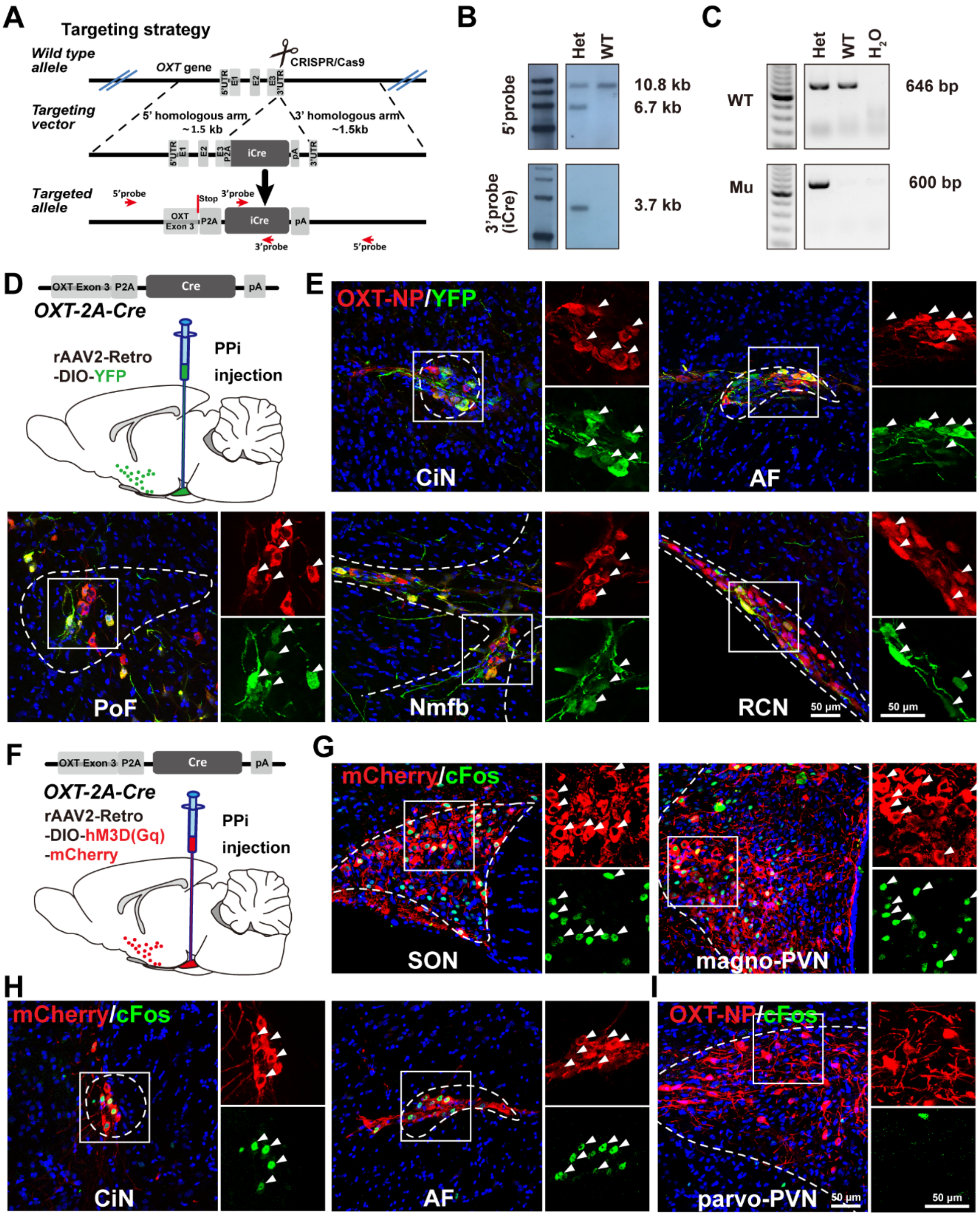

A
Targeting strategy
Wild type allele
OXT gene
CRISPR/Cas9
Targeting vector
5' homologous arm ~1.5 kb
3' homologous arm ~1.5kb
iCre
Targeted allele
5'probe
Stop
3'probe
OXT Exon 3
P2A
iCre
pA
3'probe
5'probe
B
Het
WT
5'probe
10.8 kb
6.7 kb
3'probe (iCre)
3.7 kb
C
Het
WT
$H_2O$
WT
646 bp
Mu
600 bp
D
OXT Exon 3
P2A
Cre
pA
OXT-2A-Cre
rAAV2-Retro -DIO-YFP
PPi injection
E
OXT-NP/YFP
CiN
AF
PoF
Nmfb
RCN
50 µm
50 µm
F
OXT Exon 3
P2A
Cre
pA
OXT-2A-Cre
rAAV2-Retro -DIO-hM3D(Gq) -mCherry
PPi injection
G
mCherry/cFos
SON
magno-PVN
H
mCherry/cFos
CiN
AF
I
OXT-NP/cFos
parvo-PVN
50 µm
50 µm


**Figure S5**

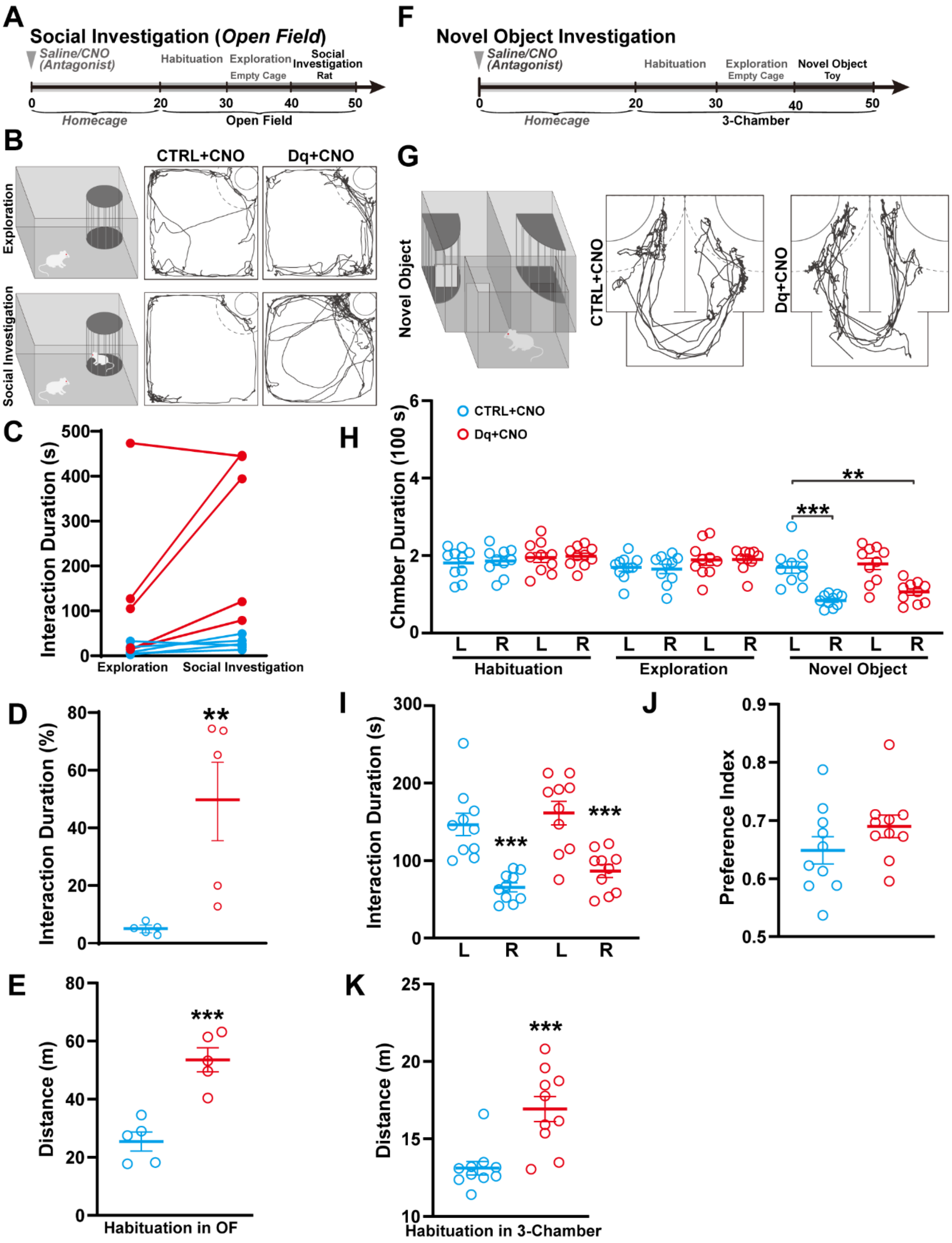
A
Social Investigation (Open Field)
Saline/CNO (Antagonist)
Habituation
Exploration
Empty Cage
Social Investigation
Rat
0
20
30
40
50
Homecage
Open Field
B
CTRL+CNO
Dq+CNO
Exploration
Social Investigation
C
Interaction Duration (s)
500
400
300
200
100
0
Exploration
Social Investigation
D
Interaction Duration (%)
80
60
40
20
0
**
E
Distance (m)
80
60
40
20
0
***
Habituation in OF
F
Novel Object Investigation
Saline/CNO (Antagonist)
Habituation
Exploration
Empty Cage
Novel Object
Toy
0
20
30
40
50
Homecage
3-Chamber
G
Novel Object
CTRL+CNO
Dq+CNO
H
CTRL+CNO
Dq+CNO
Chmber Duration (100 s)
6
4
2
0
**
***
L
R
L
R
L
R
L
R
L
R
L
R
Habituation
Exploration
Novel Object
I
Interaction Duration (s)
300
200
100
0
***
***
L
R
L
R
J
Preference Index
0.9
0.8
0.7
0.6
0.5
K
Distance (m)
25
20
15
10
***
Habituation in 3-Chamber

**Figure S6**
